# Synaptic plasticity is predicted by spatiotemporal firing rate patterns and robust to *in vivo*-like variability

**DOI:** 10.1101/2021.08.03.454974

**Authors:** Daniel B. Dorman, Kim T. Blackwell

## Abstract

Synaptic plasticity, the experience-induced change in connections between neurons, underlies learning and memory in the brain. Most of our understanding of synaptic plasticity derives from *in vitro* experiments with precisely repeated stimulus patterns; however, neurons exhibit significant variability *in vivo* during repeated experiences. Further, the spatial pattern of synaptic inputs to the dendritic tree influences synaptic plasticity, yet is not considered in most synaptic plasticity rules. Here, we address the sensitivity of plasticity to trial-to-trial variability and delineate how spatiotemporal synaptic input patterns produce plasticity with *in vivo*-like conditions using a data-driven computational model with a calcium-based plasticity rule. Using *in vivo* spike train recordings as inputs, we show that plasticity is strongly robust to trial-to-trial variability of spike timing, and derive general synaptic plasticity rules describing how spatiotemporal patterns of synaptic inputs control the magnitude and direction of plasticity. Specifically, a high temporal input firing rate to a synapse late in a trial correlated with neighboring synaptic activity produces potentiation, while an earlier, moderate firing rate that is negatively correlated with neighboring synaptic activity produces depression. Together, our results reveal that calcium dynamics can unify diverse plasticity rules and reveal how spatiotemporal firing rate patterns control synaptic plasticity.

## Introduction

Synaptic plasticity—the activity-dependent modification of synaptic strength—is widely hypothesized as the neural substrate of learning and memory throughout the brain (***Takeuchi et al., 2014***). For instance, synaptic plasticity in mammalian striatum (***Perrin and Venance, 2019***), cortex (***Buonomano and Merzenich, 1998***), hippocampus (***Martin and Morris, 2002***), and amygdala (***Bocchio et al., 2017***) have been linked to procedural, sensorimotor, associative, and emotional learning and memory, respectively. Learning requires that repeated experiences produce a stable, persistent change in synaptic connections which in turn produce stable neural activity and behavioral responses (***Abraham and Robins, 2005; Josselyn and Tonegawa, 2020***). *In vivo* experiments have revealed changes in synaptic strength, and generation, elimination, growth or shrinkage of dendritic spines (sites of synaptic input) (***Fisher et al., 2017; Trachtenberg et al., 2002; Winnubst et al., 2015; Zhang et al., 2015***). However, evidence for stable synaptic changes in response to repeated stimuli primarily comes from *in vitro* brain-slice experiments with precisely repeated input stimuli patterns, which reveal that stimulus timing, frequency, and synaptic location can control development of long term potentiation (LTP) or long term depression (LTD) (***Sjöström et al., 2001; van Rossum et al., 2000; Caporale and Dan, 2008; Lovinger et al., 1993; Hawes et al., 2013***). Yet, it is unclear if *in vitro* plasticity discoveries that used precise stimulation patterns are reproducible in the highly variable neural activity conditions *in vivo* during natural learning and behavior. Indeed, one of the great unsolved questions in neuroscience is whether stable, long-lasting synaptic plasticity occurs *in vivo* given variable neural activity, and, if so, how robust is synaptic plasticity to variability in response to repeated sensory stimuli and behaviors?

Variability and noise are prominent throughout the brain (***Faisal et al., 2008***). For instance, cortical neurons exhibit significant trial-to-trial variability *in vivo* in response to the same repeated external sensory stimulus (***Shadlen and Newsome, 1998; Stevens and Zador, 1998***). Trial-to-trial variability includes variance in the timing of individual spikes, as well as variance in firing rate over time. Thus, a given postsynaptic neuron could experience variability in the timing and frequency at each of its thousands of synaptic inputs, which would together produce (in addition to cell-intrinsic sources of variability) highly variable output spiking of the postsynaptic neuron. For a postsynaptic neuron to become potentiated or depressed in response to a specific stimulus represented by a subset of its synaptic inputs, plasticity must be robust to variance in the signal (spiking of that particular subset of inputs) as well as variance in the noise (the other synaptic inputs not related to a stimulus). In the *in vitro* case, not only is presynaptic and postsynaptic spike timing much less variable during repetition, but also the spatial pattern on the dendritic tree of the repeatedly activated synapses is likely less variable *in vitro* as well.

Experiments suggest that spatial organization of synaptic inputs on the dendritic tree are important for plasticity. For instance, *in vitro*, cortical synaptic plasticity has been shown to depend not only on rate and timing, but also on cooperativity of inputs (***Sjöström et al., 2001***), yet it is unclear how *in vivo-like* variability may affect each of these factors. Additionally, the spatial organization of synaptic inputs on the dendritic tree can affect plasticity by cooperativity among clustered inputs *in vitro* (***Brandalise et al., 2016; Kastellakis et al., 2015; Sjöström et al., 2008; Weber et al., 2016; Golding et al., 2002***). Yet, the effect of variability on spatial cooperativity under *in vivo*-like conditions is unclear, as trial-to-trial variability may affect different spatial patterns, potentially producing cooperative plasticity one trial but not another.

Intracellular calcium elevation is required for most forms of synaptic plasticity, and the spatiotemporal dynamics of the calcium signal—its peak, duration, and location—can determine the occurrence and direction (potentiation or depression) of plasticity (***Evans and Blackwell, 2015; Nevian and Sakmann, 2006; Zucker, 1999***). In spiny projection neurons (SPNs) of the striatum—input nucleus of the basal ganglia and the focus of this paper—calcium signaling may provide an eligibility trace for corticostriatal plasticity (either LTP or LTD) underlying reinforcement learning for goal-directed or habitual behavior (***He et al., 2015; Kreitzer and Malenka, 2008***). We have previously shown that synaptic calcium transients are highly sensitive to spatiotemporal patterns of synaptic input and can encode synapse-specificity and cooperativity of neighboring synaptic activity in an experimentally-validated biophysical SPN model with dendritic branches (***Dorman et al., 2018***). We have also shown that a calcium based plasticity rule with dual amplitude- and duration-thresholds can predict LTP and LTD outcomes of several in vitro plasticity experiments (***Jędrzejewska-Szmek et al., 2017***). Together, our prior works suggests that our calcium based plasticity rule, implemented in biophysical models with realistic morphology, could predict whether plasticity is robust to spatiotemporal trial-to-trial variability and which specific spatiotemporal patterns produce LTP or LTD.

Here, we investigate computationally the effect of spatiotemporal synaptic activity patterns and trial-to-trial variability to predict if persistent synaptic plasticity occurs with *in vivo*-like synaptic variability. We find that persistent plasticity is robust to trial-to-trial variability, and we also demonstrate that the spatial pattern of activity on the dendritic branch is critical for determining whether LTP or LTD occurs.

## Results

### Data-driven SPN model exhibits calcium-based synaptic plasticity for *in vivo*-like inputs

To investigate the effects of *in vivo*-like corticostriatal synaptic inputs on corticostriatal plasticity, we created a realistic biophysical SPN model (***Prager et al., 2020***) with a calcium-based plasticity rule (***Jędrzejewska-Szmek et al., 2017***) and simulated *in vivo*-like synaptic input patterns. The multi-compartment, multi-ion channel model was optimized to fit electrophysiological data using an extended version of a parameter optimization algorithm we developed (***Jędrzejewski-Szmek et al., 2018; Dorman and Blackwell, 2021***). The model reproduced the characteristic electrophysiological responses of SPNs.

Trial-to-trial variability is regularly observed in *in vivo* spike train recordings but the effect of variability on plasticity is unclear. To simulate synaptic plasticity in response to trial-to-trial variability, we obtained spike train recordings from the anterior lateral motor cortex from a published dataset (***Li et al., 2015***) from which we constructed synaptic inputs to the model. An initial single trial of corticostriatal inputs was constructed from all the spike trains of 22 behaviorally similar experimental trials to generate sufficient synaptic drive while maintaining, as much as possible, potential within-trial correlations between neurons present in the dataset. The initial one-second trial (shown as raster plot and peri-stimulus time histogram in Figure 1A,B) produced depolarization and spiking in the SPN (Figure 1C) with a firing rate consistent with *in vivo* observations.

**Figure 1.**
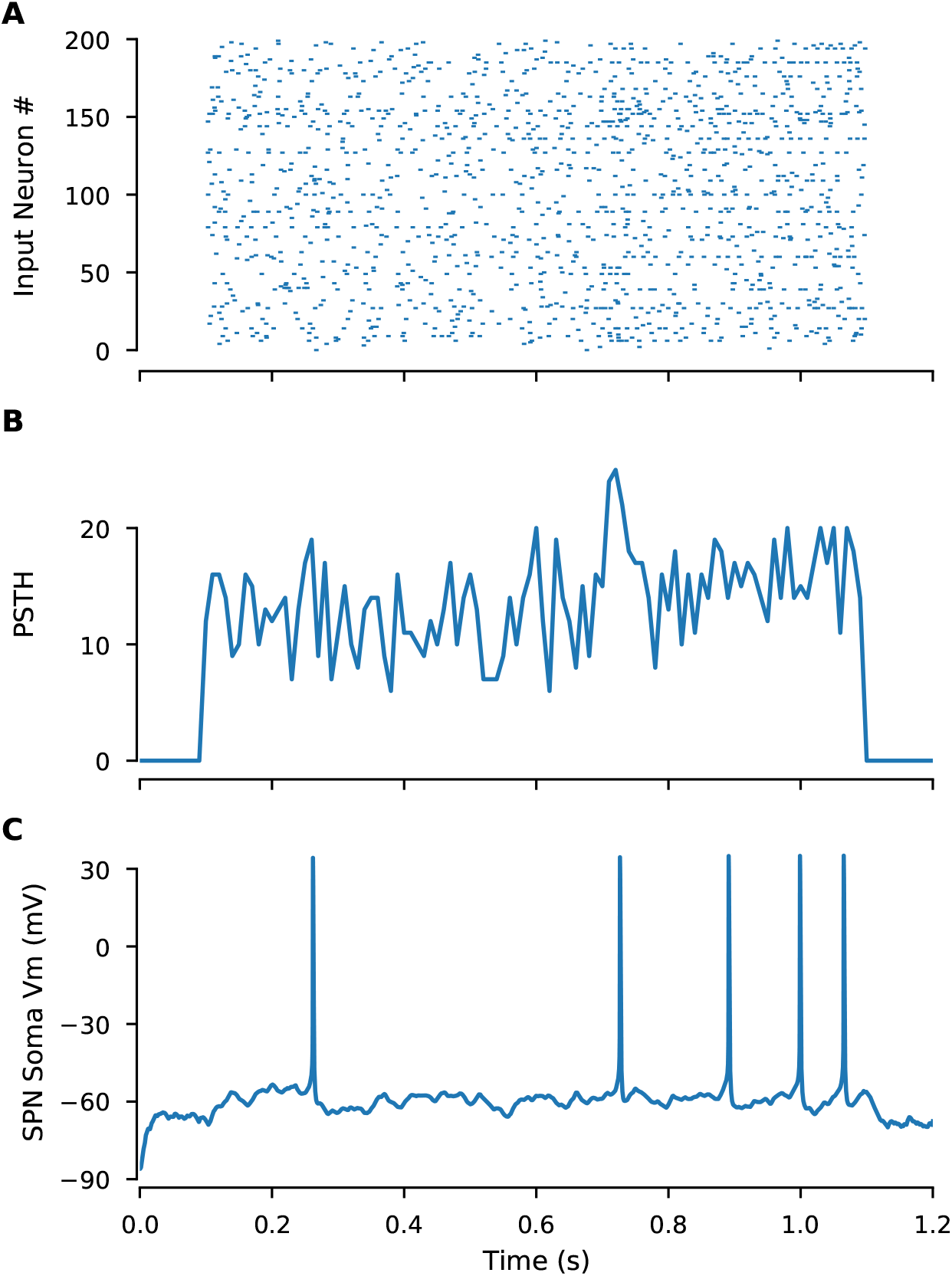
*In vivo*-like inputs constructed from cortical spike trains produce spiking in SPNs. **A**. Raster plot shows spike times for each cortical input in the model, constructed from in vivo spike train recordings. **B**. Peri-stimulus time histogram of the above raster plot (spike counts per 10 ms bin) **C**. Somatic membrane potential of the SPN model showing spiking output induced by cortical input.

The first question addressed is whether a calcium-based synaptic plasticity rule that was derived to explain STDP data is sufficiently general to produce synaptic plasticity in response to spatiotemporally distributed synaptic inputs. To determine whether this calcium-based plasticity rule would predict plasticity for *in vivo* like conditions, we simulated synaptic weight changes in response to trial-to-trial variability. We used our calcium-based plasticity rule that can reproduce results from several spike-timing dependent plasticity experiments on SPNs *in vitro* (***Jędrzejewska-Szmek et al., 2017***). Crucially, this rule is entirely based on spine calcium dynamics, not relative spike timings, so it is a general rule that encompasses both frequency-based and spike-timing plasticity rules.

Cortical spike trains indeed produced synaptic plasticity with our calcium-based plasticity rule. We examined the spine calcium concentration and the synaptic weight of every synapse in response to a single 1-second trial of randomly distributed *in vivo* spike trains. We found that, at the end of a single trial, some synapses exhibited potentiation, some exhibited depression, and others exhibited no change (Figure 2—example synapses; Figure 3—all synapses), with most exhibiting little change. These results show that a calcium-based plasticity rule determined from *in vitro* data can produce plasticity with spatially distributed *in vivo*-like synaptic input conditions.

**Figure 2.**
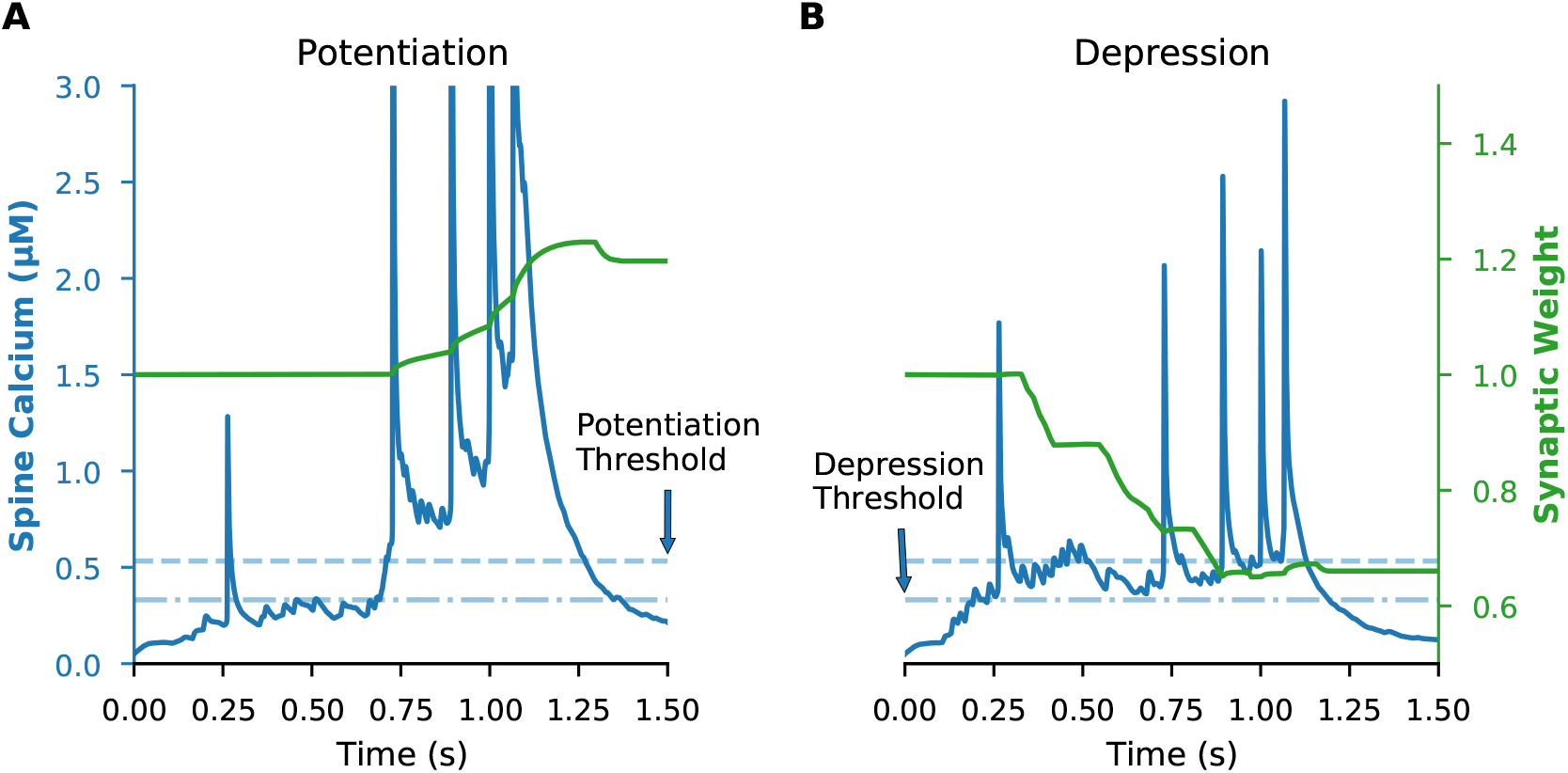
Calcium-based plasticity rule produces both potentiation and depression. A calcium-based plasticity rule was implemented with dual amplitude and duration thresholds. LTD required that spine calcium concentration exceed the amplitude threshold (dot-dashed line) of 0.33 μM for greater than 28 ms, while LTP required that spine calcium concentration exceed a higher amplitude threshold (dashed line) of 0.53 μM for at least 3.3 ms. Example traces are shown of a synapse that potentiates (**A**) or depresses (**B**) following a single trial. Blue lines show spine calcium concentration (with left y-axis) and green lines show synaptic weight (with right y-axis) for each example synapse.

**Figure 3.**
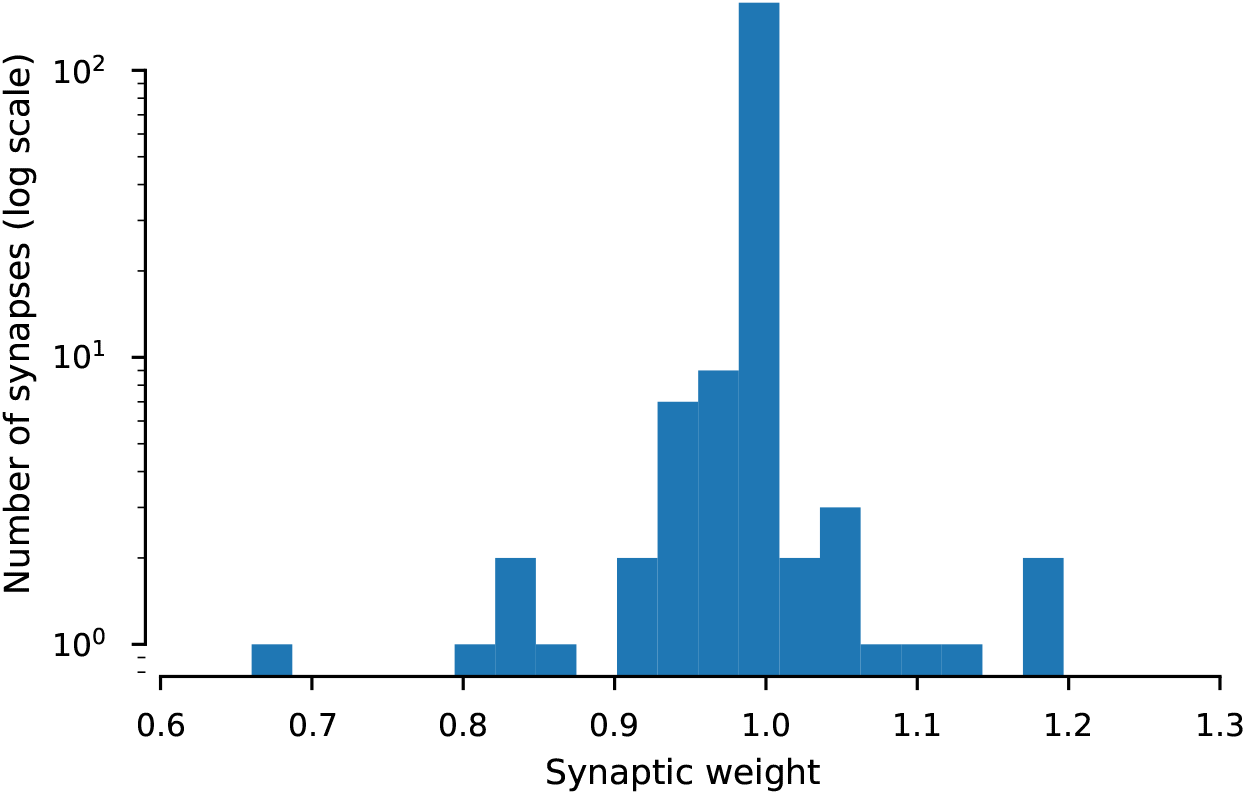
Distribution of synaptic weights after initial trial. Histogram shows the distribution of synaptic weights for all synapses following a single trial (note log scale of y axis). All weights were initialized at 1, and post-trial weights greater than 1 are potentiation while weights less than 1 are depression.

### Synaptic plasticity is highly robust to trial-to-trial variability

During repeated behaviors, cortical neurons exhibit significant levels of trial-to-trial variability, yet it remains unclear how this variability affects synaptic plasticity. For synaptic plasticity to serve as the basis of learning, it should be robust to naturally observed variability in neuron spiking. However, many plasticity experiments use highly regular, precisely repeated stimulus patterns. To bridge the gap between *in vitro* plasticity findings due to precisely repeated stimuli and *in vivo* plasticity with spatiotemporally dispersed inputs and trial-to-trial variability, we simulated the response to 10 repeated trials, with varying levels of trial-trial variability. This is analogous to 10 behavioral learning trials during which striatal neurons receive variable cortical input from trial to trial.

For every level of trial-to-trial variability we simulated, a subset of synapses exhibited robust weight change at the end of 10 repeated trials. As shown in Figure 4A, synaptic weight over time consistently accumulates potentiation or depression for a subset of synapses regardless of variability level. This robust weight change also is observed when variability is introduced by randomly moving spikes to different trains (Figure 4— supplementary figure 1). These results predict that synaptic weight change is robust to high levels of trial-to-trial spike time variability, suggesting that *in vivo* variability does support synaptic plasticity.

**Figure 4.**
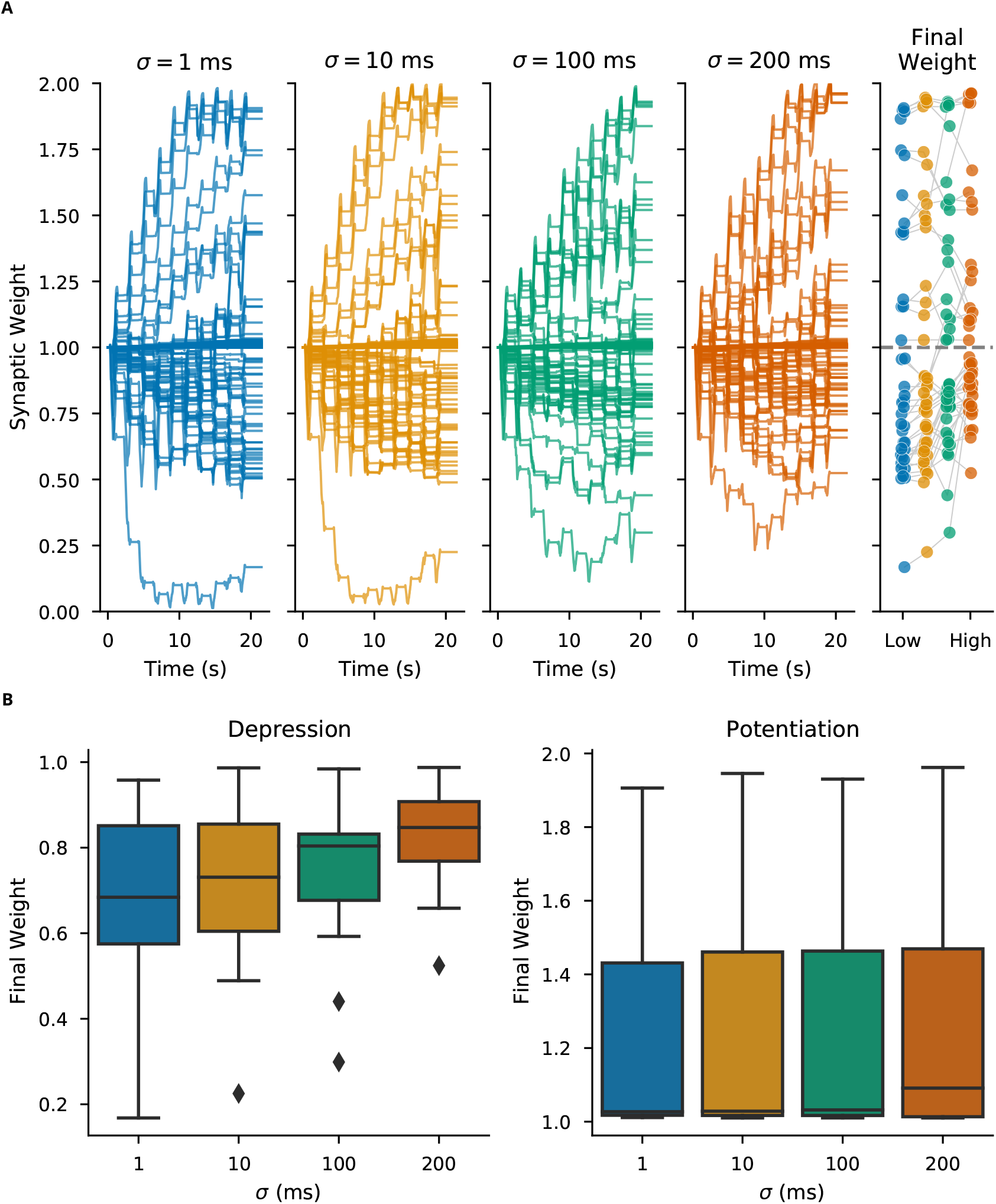
Synaptic plasticity is robust to trial-to-trial variability. (**A**) The weight of every synapse over a full 10-trial experiment is shown for different trial-to-trial variability conditions, with the sigma value corresponding to the standard deviation of random jitter of spike times. (Right) Ending weight of each synapse is shown for each variability condition (synapses with near zero weight change excluded for visualization). (**B**) Distribution of final weights grouped by potentiation and depression shows that variability reduces synaptic depression magnitudes, but has little effects on the distribution of potentiation magnitudes. Correlation of ending synaptic weight versus variability was significant for depressing synapses (R=0.306, p=0.0006, N=121 events), but not for potentiating synapses (R=0.054, p=0.510, N=148 events)

Though synaptic weight change persisted across levels of trial-to-trial variability, the magnitude of weight change at the end of an experiment was reduced for increasing levels of variability. Figure 4B shows ending synaptic weight as a function of trial-to-trial variability, demonstrating that high variability reduces the magnitude of depression, but not potentiation, both for jittered and moved spikes (Figure 4—supplementary figure 2). These results suggest that variability in spike timing primarily effects the magnitude of plasticity but rarely the direction (potentiation or depression).

### Plasticity of a single synapse is only partially predicted by its presynaptic activity

Individual synapses receive wide ranges of presynaptic input patterns and exhibit a broad range of synaptic plasticity outcomes. Which properties of synaptic input patterns, that are potentially modulated by trial-to-trial variability, predict the magnitude and direction of synaptic plasticity? To investigate this question, we first asked whether the total presynaptic spike count for a given synapse predicted that synapse’s weight at the end of 10 trials (ending weight). We compared ending weight of every synapse to its total presynaptic spike count across all 10 trials for experiments at different levels of trial-to-trial variability (Figure 5). We found a general pattern that synapses with low presynaptic spike counts exhibited little to no synaptic weight change; synapses with a moderate presynaptic spike count tended to exhibit depression; and synapses with a high presynaptic spike count exhibited potentiation. However, a wide range of ending synaptic weights was observed for synapses with moderate presynaptic spike counts, and further, these synapses were most affected by trial-to-trial variability. The synaptic weight was not correlated with the synapse’s distance to the soma, except for the lower levels of variability (Figure 5—Supplementary Figure 1). These results suggest that while spike count alone is a significant factor in predicting synaptic weight change, other spatiotemporal factors may also be important, and these factors may be significantly affected by trial-to-trial variability.

**Figure 5.**
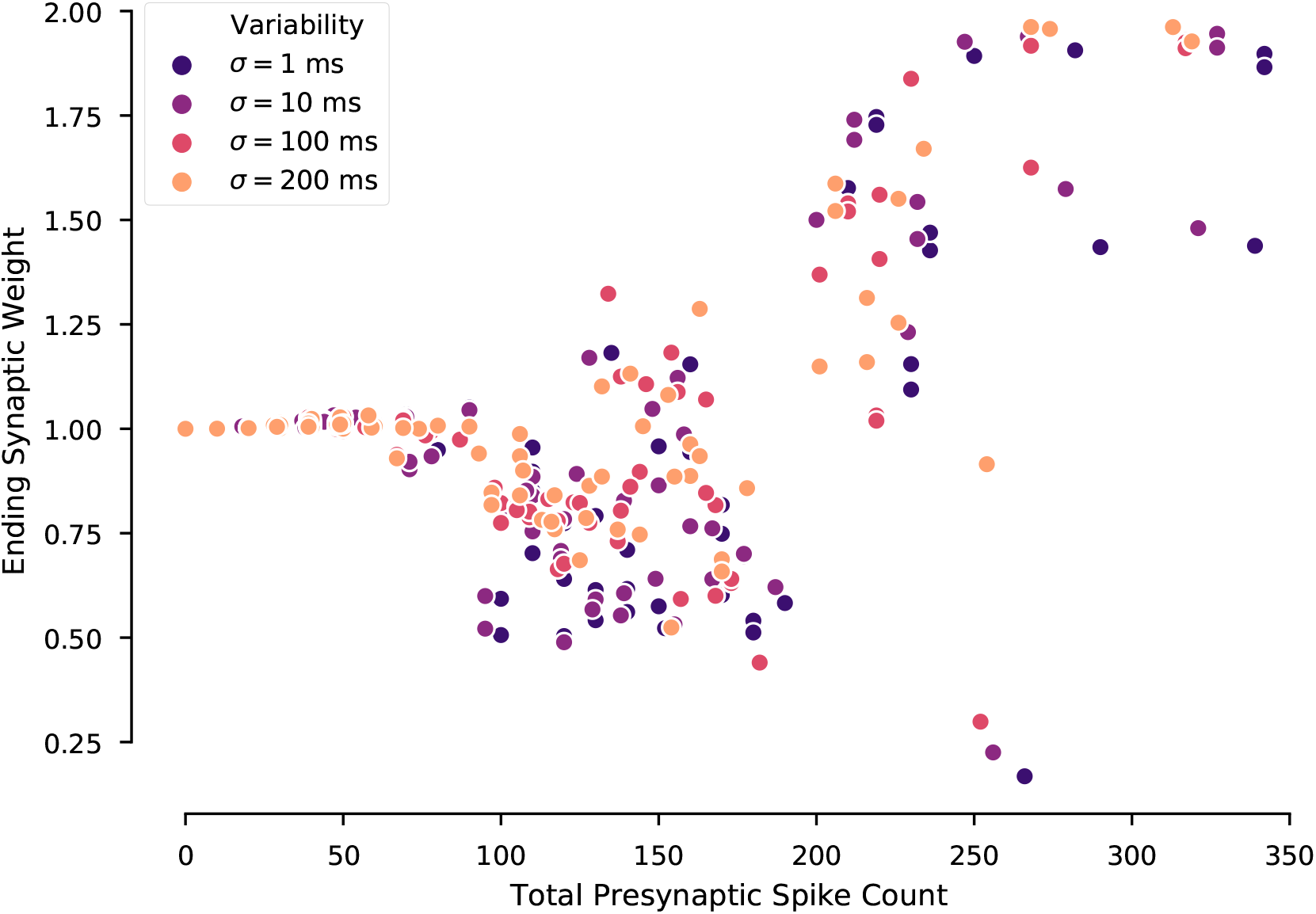
Ending synaptic weight is partially predicted by total presynaptic spike count per synapse. Ending synaptic weight of each synapse is plotted versus its total presynaptic spike count across all 10 repeated trials, with experiments separated by the level of trial-to-trial variability. Ending weight exhibits no change for low spike counts, tends toward depression for intermediate spike counts, and exhibits potentiation for high spike counts. This trend is consistent regardless of trial-to-trial variability; however, for intermediate spike counts the ending weight is highly variable.

### Plasticity of a single synapse is affected by presynaptic temporal firing rate pattern

As total presynaptic spike count alone did not completely predict ending synaptic weight per synapse, we next investigated the effect of the temporal pattern of presynaptic firing rate on synaptic plasticity. Though spike count per synapse was consistent for experiments, the trial-to-trial variability introduced changes in spike timing that altered the time-varying instantaneous presynaptic firing rate for each synapse. To identify whether instantaneous firing rate over the course of a single trial was associated with potentiation, depression, or no-change, we computed a weight-change triggered average presynaptic firing rate. This weight-change triggered average was computed by binning trials and synapses based on the magnitude of synaptic weight change following an individual trial, computing the instantaneous presynaptic firing rate vs. time for each synapse and trial, and averaging across synapse-trials within each bin.

Our results show that synapses that strongly potentiate exhibit a weight-change-triggered average presynaptic firing rate with a high peak firing rate late in the trial. In contrast, synapses that strongly depress exhibit an earlier peak firing rate or a moderate sustained presynaptic firing rate. Synapses with little or no weight change exhibit a low presynaptic firing rate (Figure 6A). This pattern was also observed when variability was introduced by moving spikes between trains (Figure 6—supplementary figure 1A). We also calculated the weight-change-triggered average calcium concentration, to assess how presynaptic firing dynamics were translated into calcium elevations. As shown in Figure 6B (and Figure 6—supplementary figure 1B), synapses that potentiate have higher calcium concentration than synapses that depress, and synapses that strongly potentiate have the highest calcium. Though the largest differences are in the second half of the trial, weight change dependent differences in calcium concentration for synapses that depress are more apparent in the first half of the trial. These results suggest that input timing within a trial, which affects peak instantaneous firing rate and calcium concentration, is a critical factor for the direction and amplitude of synaptic weight change. Though temporal pattern discriminates strong LTP from strong LTD, the temporal patterns for moderate plasticity are not as clear. Thus, next we evaluated the role of spatial patterns of synaptic input.

**Figure 6.**
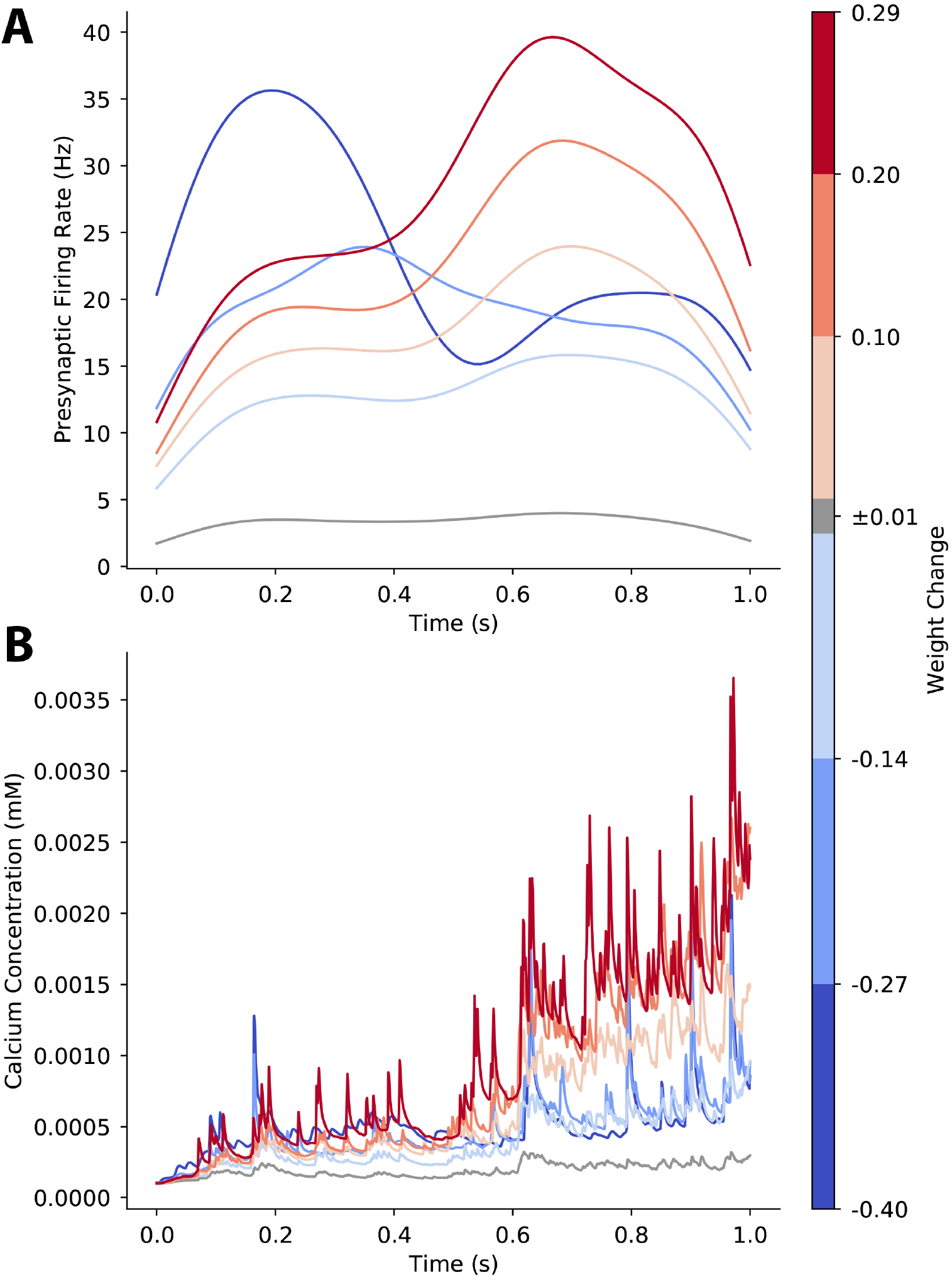
Temporal pattern determines direction of plasticity as shown by weight-change-triggered-average. **A.** For each synapse on each trial, we computed the instantaneous firing rate of its presynaptic input activity and grouped synapse-trials into bins based on the size of the synaptic weight change that occurred following a single trial. Then, we averaged across the instantaneous firing rate of each bin. Late and high peak firing rates lead to LTP, while earlier peak firing rate or moderate firing rate leads to LTD. **B.** For each synapse on each trial, we computed the calcium concentration for each synapse-trial and averaged across the calcium concentration for each weight change bin. Calcium concentration is higher during the second part of the trial, both for potentiating and depressing synapses.

### Plasticity of a single synapse is affected by neighboring synaptic activity

Prior work has shown that synaptic plasticity can be affected by spatiotemporally cooperative synaptic activity—that is, multiple synapses on the same dendritic branch active within a limited time window (***Govindarajan et al., 2011; Legenstein and Maass, 2011; Cichon and Gan, 2015; Brandalise et al., 2016; Magó et al., 2020; Weber et al., 2016; Losonczy et al., 2008***). Our prior work has shown that spatiotemporal activity patterns have nonlinear, spatially specific effects on calcium transients in dendrites and spines (***Dorman et al., 2018***). Thus, it is likely that nearby synaptic activity can cooperatively influence plasticity in our calcium-based model. As shown in figure 5, a given synapse’s presynaptic firing rate, alone, is not sufficient to fully determine its synaptic weight change. Therefore, we next investigated the cooperative effect of neighboring synaptic activity on weight change at each synapse.

We found that synapses which strongly potentiate experience higher neighboring synaptic activity in the dendritic branch than synapses which strongly depress (Figure 7A) for both types of spike train variability (Figure 7—supplementary figure 1). This suggests that cooperative synaptic activity among neighboring synapses within a dendritic branch influences the outcome of plasticity. We averaged across neighboring synapses because no spatial pattern was apparent when considering input to neighboring synapses individually (Figure 7—supplementary figure 2). To further investigate a relationship between direct and neighboring inputs for plasticity, we computed correlation coefficients between each synapse’s direct instantaneous firing rate and the combined rate of its neighbors, again binned by plasticity outcome. As shown in Figure 7B, strongly potentiating synapses were more likely to be positively correlated with neighboring activity, while strongly depressing synapses were more likely to be negatively correlated. Together, our results show that both temporal and spatial patterns are critical to synaptic plasticity.

**Figure 7.**
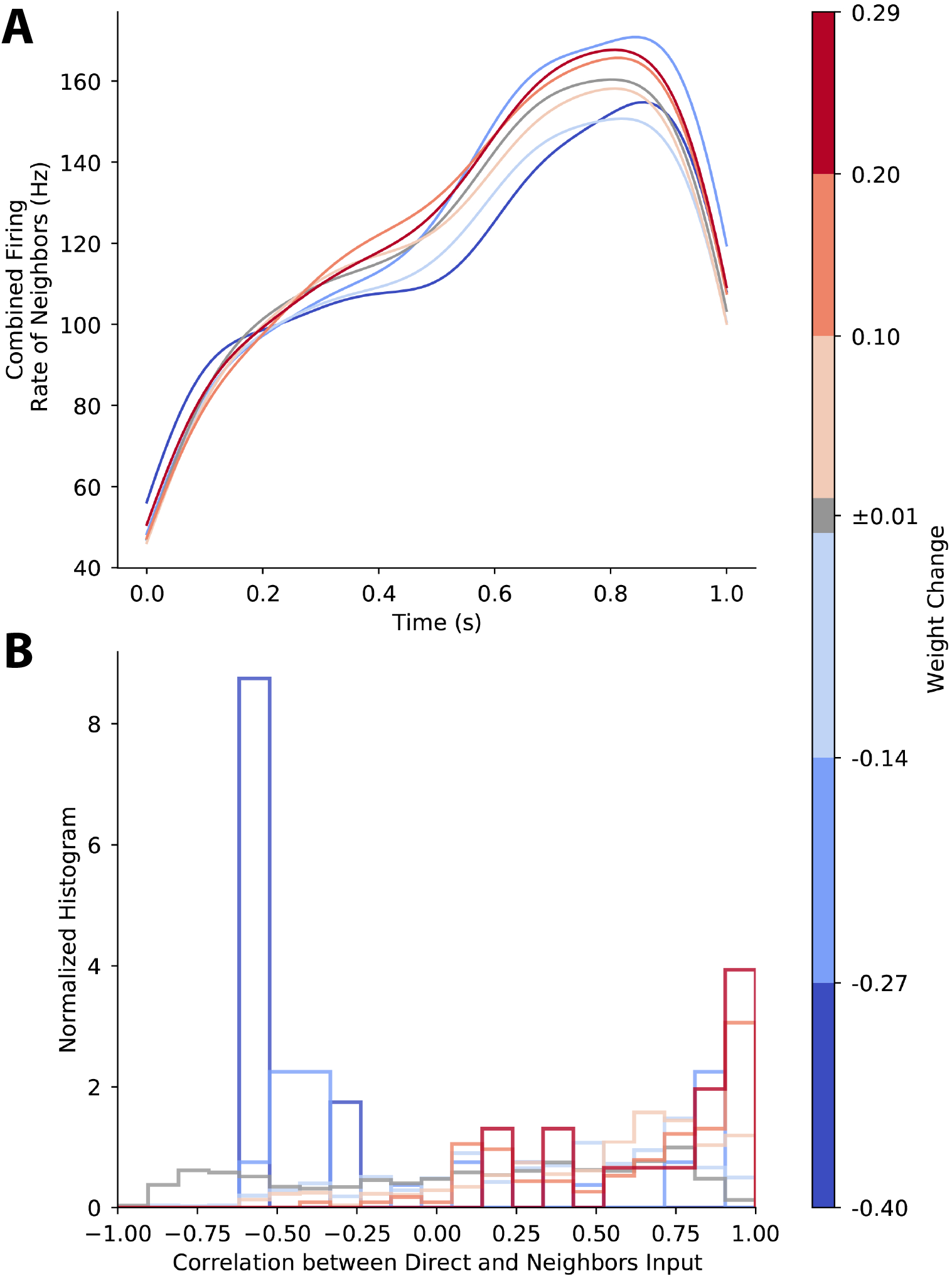
Activity of neighboring synapses influences direction of plasticity. **A.** Neighboring synaptic activity is associated with small difference in strong LTP and LTD. Traces show the combined neighboring synaptic instantaneous firing rate averaged for bins of synaptic weight change following a trial. For every synapse, its 20 nearest neighbors’ spike trains were combined and an instantaneous firing rate was computed from the combined spike train. Synapses were then binned by amount of weight change following a trial, and the average neighboring instantaneous rate was computed for each bin. **B.** Correlation between direct and neighboring synapses influences direction of plasticity. Histograms show the correlation between direct and neighboring presynaptic firing rates binned by plasticity outcome. High positive correlation between direct and neighboring firing rate is associated with LTP, while low correlations are associated with LTD.

### Dynamics of pre-synaptic firing rate predicts synaptic plasticity

To further assess robustness of the results, we repeated simulations using 1000 different variations of the mapping from spike trains to synapses. Since the change in plasticity is consistent with repeated trials (Fig 4), we simulated a single trial. The weight-change-triggered triggered pre-synaptic firing rate (Figure 7— supplementary figure 3A) reveals that, similar to Fig 6, synapses that strongly potentiate have a transiently high firing rate, though the time of peak firing varies. Regardless of the dynamics of the pre-synaptic firing rate, the calcium concentration (Figure 7— supplementary figure 3B) exhibits a higher concentration later in the trial, as observed in Fig 6B. The firing rate of neighboring synapses (Figure 7— supplementary figure 3C) reveals a much higher firing frequency for neighbors of potentiating synapses than neighbors of depressing synapses. For synapses that exhibited a weight change, the correlation between neighboring synapse firing rate and weight change was 0.1, averaged across 5 sets of simulations. Though the correlation is low, it is highly significant (P<0.0001) demonstrating that spatiotemporal firing patterns are key to determining synaptic plasticity.

To quantitatively assess how spatiotemporal pattern of synaptic input controls synaptic plasticity, we used random forest regression to predict synaptic weight change from instantaneous firing rate to the synapse, together with either firing rate of neighboring synapses, correlation between direct input and input to neighboring synapses, or distance of synapse to the soma. The instantaneous firing rate was discretized into a small number of time samples to coarsely represent the mean firing rate over time: e.g., 1 sample measures the mean firing rate for the entire trial. Figure 8 shows that increasing the resolution of sampling the firing rate improved the prediction of weight change, demonstrating that temporal pattern is important. Unexpectedly, spatial information did not improve the prediction. The same pattern is observed when variability was introduced by moving spikes between trains (Figure 8— supplementary figure 1A) or using alternative mappings of spike trains to synapses (Figure 8— supplementary figure 1B). In summary, the dynamics of direct input to the synapse is crucial for determining synaptic plasticity, but firing rate of neighboring synapses was not needed, despite the correlation between synaptic weight change and neighboring firing rate, likely due to ability of random forest regression to find temporal features.

**Figure 8.**
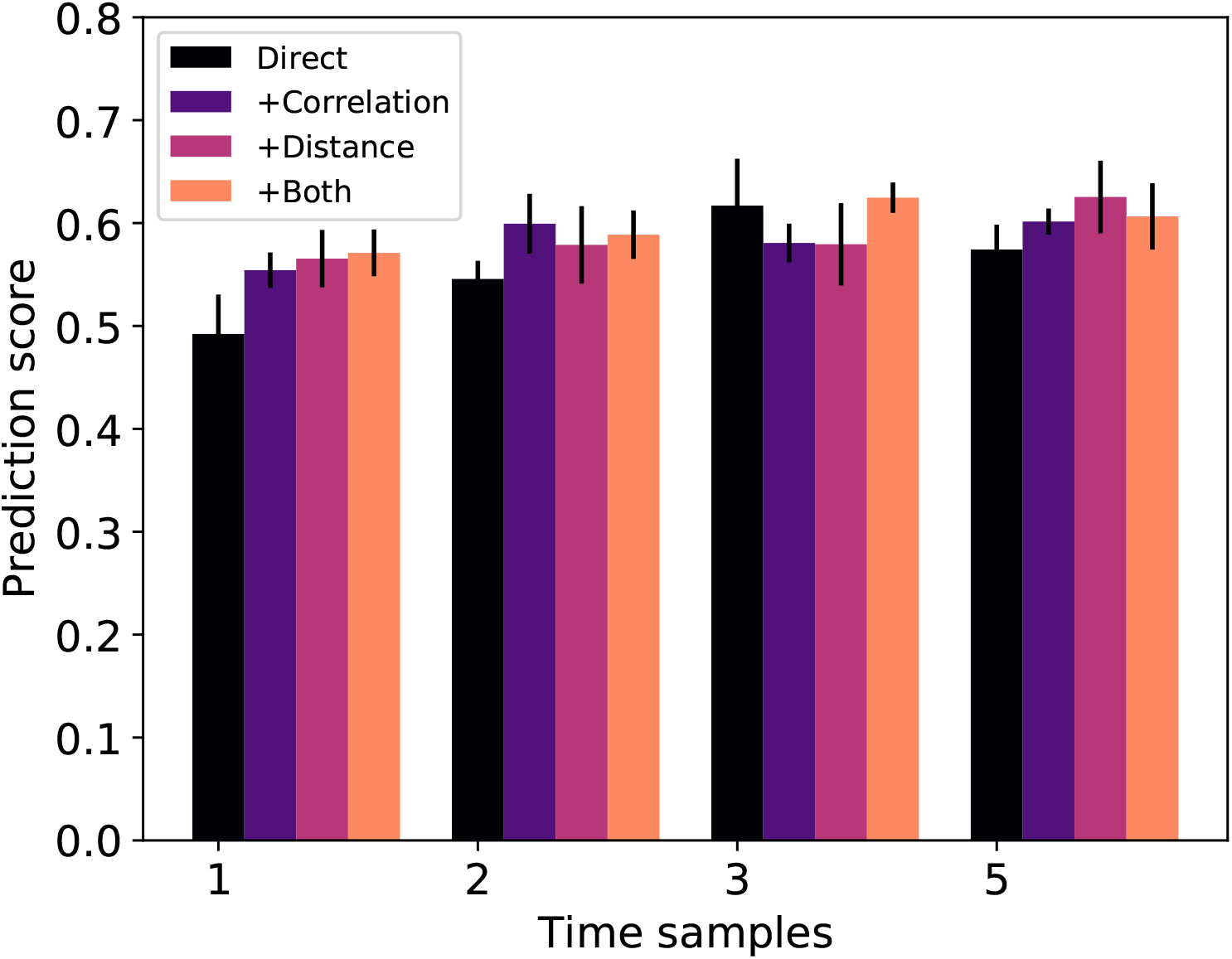
Temporal pattern of input predicts weight change better than mean firing rate. Prediction score is the coefficient of determination, R^2^, of the predicted weight change for the test set. N=4 regressions for each combination of features. Error bars show 1 standard error. ANOVA shows that increasing number of time samples improves the prediction score (F(3,76)=4.776, P=0.0042).

## Discussion

In this study, we addressed how temporal and spatial patterns of *in vivo*-like synaptic inputs with trial-to-trial variability impact the direction and magnitude of synaptic plasticity. We employed a calcium based plasticity rule in a biologically constrained computational model previously validated on *in vitro* plasticity protocols to predict plasticity for *in vivo*-like conditions. We found that synaptic plasticity with *in vivo*-like activity is robust to trial-to-trial variability. Further, we found that the temporal pattern of synaptic inputs within a trial and the spatial pattern of neighboring synaptic inputs on the dendritic tree controls synaptic plasticity outcomes, with a transient high firing frequency producing strong LTP and a moderate increase in firing frequency producing LTD. The firing frequency of neighboring synapses relates to the plasticity outcome, with correlation or high firing frequency of neighbors producing LTP and lack of correlation or low firing frequency of neighbors producing LTD. Our work provides key insights into the nature of synaptic plasticity in conditions more likely to occur during natural behavior. Together, these results predict that plasticity is highly robust to variable spike timing in the brain and suggest that both temporal and spatial aspects of synaptic integration are critical to plasticity.

Our finding that plasticity at a single synapse is influenced by neighboring synaptic activity is consistent with many studies (both *in vivo* and *in vitro*) showing relationships between spatial synaptic input patterns and plasticity. For instance, in SPNs *in vitro*, spatially clustered synaptic inputs can produce supralinear depolarization (termed NMDA-spikes or plateau potentials) and synaptic calcium transients (***Plotkin et al., 2011; Dorman et al., 2018; Prager et al., 2020; Du et al., 2017***), and synaptic activation of 2-4 neighboring spines at depolarized potentials can produce nonlinear enhancement of spine calcium transients (***Carter et al., 2007***). Building on these findings, our work predicts that neighboring synaptic interactions can influence the direction and magnitude of corticostriatal synaptic plasticity *in vivo*, with high neighboring activity producing LTP and low neighboring activity producing LTD. Spatially clustered inputs have been shown to induce synaptic potentiation (termed cooperative LTP) in cortical and hippocampal pyramidal neurons *in vitro* (***Brandalise et al., 2016; Golding et al., 2002; Gordon et al., 2006; Larkum et al., 2009; Losonczy et al., 2008; Makara and Magee, 2013; Schiller et al., 2000; Weber et al., 2016; Magó et al., 2020***). Similarities in synaptic integration properties of SPNs and pyramidal neurons (***Oikonomou et al., 2014***) suggest that our calcium-based plasticity rule could be implemented in models of pyramidal neurons and could account for cooperative LTP observations *in vitro* and predict *in vivo* plasticity in pyramidal neurons.

Observations suggest that functional synaptic clustering is a key component in plasticity and learning. For instance, correlated activity in spatially clustered spines has been observed *in vivo* in pyramidal neurons (***Takahashi et al., 2012; Winnubst et al., 2015; Wilson et al., 2016; Kerlin et al., 2018***). Further, *in vivo* calcium transients in neighboring spines of the same dendritic branch correlate with structural potentiation of spines and behavioral learning in motor cortical neurons (***Cichon and Gan, 2015***). This is consistent with our observation that strongly potentiating synapses had higher synaptic inputs than depressing synapses. However, it is unclear whether functional clustering arises from synaptic connectivity in development, or if activity dependent plasticity can generate functional clusters starting from random connectivity. Our results, which used randomly distributed inputs, suggest that spatial synaptic correlations may emerge with random spatial distribution of inputs, thereby potentiating neighboring synapses that each receive higher than average input and producing functional synaptic clusters. These implications of our results are consistent with another modeling study investigating the impact of synaptic clustering on somatic membrane potential with *in vivo*-like conditions (***Ujfalussy and Makara, 2020***), which suggested that global plasticity rules would not be sufficient for formation of synaptic clusters. ***Ujfalussy and Makara (2020)*** predicted that local plasticity rules would be necessary, though their model did not implement local plasticity rules to test that prediction. Our calcium-based plasticity rule, which accounts for both local and global plasticity effects, is a method for formation of synaptic clusters, consistent with ***Ujfalussy and Makara (2020)*** as well as experimental evidence. The resulting synaptic plasticity accounts for the impact of local dendritic activity, and the potentiation of correlated synapses can produce spatial clustering. Our results also suggest a role for spatial patterns in synaptic depression. Specifically, whereas high correlations among neighboring synapses were associated with potentiation, synapses that depressed had negatively correlated or lower than average neighboring synaptic activity.

This suggests a mechanism for spatially balanced potentiation and depression, which could provide homeostatic balance and prevent runaway potentiation. These implications are consistent with *in vivo* experiments that have shown spine shrinkage in inactive spines accompanying structural potentiation of nearby spines, suggesting heterosynaptic depression as an important compensatory plasticity mechanism (***Oh et al., 2015; El-Boustani et al., 2018***). Our results suggest that our calcium-based plasticity rule effectively captures the impact of neighboring synaptic activity on depression to support compensatory plasticity. Predictions from our results could be experimentally tested *in vitro* using glutamate uncaging to apply temporally correlated or uncorrelated patterns to neighboring dendritic spines, together with whole cell recording to measure LTP or LTD and calcium imaging of dendritic spines to relate stimulation patterns with calcium activity and synaptic plasticity.

The distinct temporal patterns that were associated with potentiation or depression have implications for striatal and wider basal ganglia circuit function. Though corticostriatal LTP requires dopamine in addition to calcium elevation (***Fisher et al., 2017***), prior experimental work has shown calcium-dependent synaptic eligibility traces in SPNs (***Shindou et al., 2019***). These eligibility traces for corticostriatal LTP exhibit a temporal dependence such that an LTP protocol followed within a few seconds by dopaminergic stimulation produces LTP (***Yagishita et al., 2014***). This pattern of cortical inputs followed by dopamine is consistent with reinforcement learning, as the rewarding outcome (represented in the striatal dopamine signal) temporally follows the action that produced it (represented in corticostriatal activity). Thus, though our plasticity model doesn’t account for dopamine directly, we suggest that our calcium-based plasticity rule accounts for eligibility traces for LTD or LTP and captures the spatiotemporal pattern of corticostriatal activity that, when followed by dopaminergic stimulation, produces plasticity. Further, as dopaminergic activity is consistent with volume transmission (***Borroto-Escuela et al., 2018; Zoli et al., 1998***), we suggest that our calcium-based plasticity rule provides the spatiotemporal specificity indicating which synapses are eligible for reinforcement when followed by a spatially diffuse dopamine signal.

This study has important implications for plasticity in variable *in vivo* conditions by showing that precise spike timing is not required. Our work suggests that while plasticity is sensitive to firing rate, it is highly robust to variance in precise spike timing during a trial. *In vitro* corticostriatal spike-timing dependent plasticity experiments demonstrated that NMDAR-dependent (and spike-timing dependent) LTP is highly sensitive to jittered spike timings, though increasing pairings of presynaptic and postsynaptic stimuli could recover LTP (***Cui et al., 2018***). Thus, precise spike timing rules could make *in vivo* plasticity unlikely, given variability of spike timing (***Williams et al., 2019***). Importantly, our calcium-based plasticity rule is independent of spike timing, though we previously showed it could reproduce *in vitro* spike timing dependent plasticity results. Thus, we suggest our rule is generalizable to *in vivo* conditions, and we predict that plasticity *in vivo* is robust to variable spike timing. Consistent with this implication, other work has shown that cortical and striatal neurons exhibit decreasing trial-to-trial variability *in vivo* during learning that corresponds to reduced behavioral variability of the learned action, and this reduction is dependent on striatal plasticity (***Santos et al., 2015***). Our result that potentiation is robust to high trial-to-trial variability suggests that corticostriatal plasticity may occur even with highly variable conditions early in learning and then, by potentiating the relevant synapses, produce decreased variability in striatal spiking with learning.

Our work is consistent with a prior model that predicted that plasticity was not sensitive to spike timing with *in vivo* like firing patterns (***Graupner et al., 2016***). Nonetheless, our research is a major advance over that prior work which implemented a plasticity rule based on simplified calcium dynamics in a non-spatial model (***Graupner and Brunel, 2012***) or derived firing rate plasticity models from simplified calcium dynamics (***Lappalainen et al., 2019***). Our calcium-based rule is implemented with detailed biophysical models of calcium dynamics in a neuron model that includes dendritic morphology. More importantly, we extend prior work to show that temporal patterns are still important, and that spatial interactions among synapses influence plasticity. We also predict that potentiation and depression are differentially sensitive to both spatiotemporal patterns and trial-to-trial variability.

Our results show that the spatial aspects of synaptic integration may contribute to synaptic plasticity. However, spiking network models that incorporate spike-timing dependent plasticity rules to investigate the effect of plasticity on network activity neglect spatial patterns of synaptic inputs to a single neuron, reducing neurons to point processes (***Legenstein et al., 2005; Berthet et al., 2016; Dunovan et al., 2019***). We suggest that our calcium based plasticity rule could be used in future work to develop simplified plasticity models incorporating both temporal and spatial effects of synaptic activity.

A statistical model using temporal and spatial kernels to predict the effect of both direct and neighboring synaptic activity on the synaptic weight of each synapse could be derived from biophysical single neuron models with our calcium based rule and then simulated efficiently in large networks with simplified neuron models. Incorporating this synaptic plasticity rule in large scale simulations of network models of the basal ganglia could then better predict how corticostriatal plasticity supports goal-directed and habit learning and identify potential therapeutic targets for modulating aberrant plasticity in addiction.

## Methods

We developed a biologically-constrained computational SPN model with multiple ion channels identified in SPNs, real dendritic morphology, explicit dendritic spines, sophisticated calcium dynamics, and a calcium-based plasticity rule. Model parameters were determined using our parameter optimization software to fit the model to electrophysiological current injection data (***Dorman and Blackwell, 2021***). To generate *in vivo*-like synaptic inputs, we obtained and analyzed anterior lateral motor cortical spike trains from the CRCNS.org repository. The model was simulated with these *in vivo* spike trains to investigate whether plasticity occurs with *in vivo*-like activity.

### SPN model morphology and passive membrane properties

A biophysical SPN model we previously published (***Dorman et al., 2018***) was adapted and translated from the GENESIS simulator format to our declarative format (***Blackwell et al., 2021***) for the MOOSE simulator (https://moose.ncbs.res.in). We used a D1 SPN morphology obtained from the Luebke repository (***Goodliffe et al., 2018***) on neuromorpho.org (***Ascoli et al., 2007***). Dendritic spines were modeled both implicitly and explicitly. Explicit dendritic spines were modeled for synaptic inputs and calcium dynamics at a density of 0.1 spines/μm with cylindrical head (0.5 μm diameter, 0.5 μm length) and neck (0.12 μm diameter, 0.5 μm length) on dendritic branches greater than 25 μm from the soma. The density of explicitly modeled spines is not representative of the full spine density observed experimentally, and modeling the full spine density would be computationally intensive. Therefore, the implicit effect of dendritic spines (those not explicitly modeled) on passive membrane properties and dendritic channel densities was modeled by compensating dendritic membrane resistivity (***RM***), membrane capacitivity (***CM***), and axial resistivity (***RA***), as well as channel maximal conductance values (***Holmes et al., 2006***) using a distance dependent function fit to experimentally observed spine density versus distance from the soma (***Wilson, 1992***). Values for ***RM, CM***, and ***RA*** were set to 6.02 Ωm^2^, 0.011 F/m^2^, and 1.3 Ωm respectively, based on automatic parameter optimization.

### Voltage gated ion channels

As previously described (***Dorman et al., 2018***), the model incorporates the following voltage gated sodium and potassium ion channels that have been observed in SPNs: a fast sodium channel (NaF) (***Ogata and Tatebayashi, 1990***); fast (Kaf/Kv4.2) (***Tkatch et al., 2000***) and slow (Kas/Kv1.2) (***Shen et al., 2004***) A-type potassium channels; an inwardly rectifying potassium channel (Kir2) (***Steephen and Manchanda, 2009***); a resistant persistent potassium channel (Krp; also called delayed rectifier) (***Nisenbaum and Wilson, 1995***); a big conductance voltage- and calcium-activated potassium channel (BK) (***Berkefeld et al., 2006***); and a small conductance calcium-activated potassium channel (SK) (***Maylie et al., 2004***). Six voltage gated calcium channels (VGCCs) are also included: R-type (CaR/Cav2.3) (***Brevi et al., 2001; Foehring et al., 2000***), N-type (CaN/Cav2.2) (***Bargas et al., 1994; Kasai and Neher, 1992; McNaughton and Randall, 1997***), two L-type (CaL1.2/Cav1.2 and CaL1.3/Cav1.3) (***Bargas et al., 1994; Kasai and Neher, 1992; Tuckwell, 2012***), and two T-type (CaT3.2/Cav3.2/α1H and CaT3.3/Cav3.3/α1I) (***McRory et al., 2001***). We newly added a calcium activated chloride channel (CaCC) based on the ANO2/TMEM16B channel, which has been observed in SPNs and allows better fit of the AHP waveform (***Song et al., 2016; Pifferi et al., 2009***). Channel kinetic equations are the same as our previously reported model, but parameters were updated during the parameter optimization by allowing half-activation voltages and time constants to be modified by the parameter optimization algorithm.

Model optimization was done using ajustador software (***Jędrzejewski-Szmek et al., 2018; Dorman and Blackwell, 2021***) that was updated to allow variation of channel kinetics, in addition to channel conductances and membrane properties, in order to fit the model response to experimental data for injected current steps. Briefly, the parameter optimization algorithm utilized a covariance matrix adaptation with evolutionary strategy (cma-es) to vary model parameters (***Hansen et al., 2019***), and a feature-based fitness metric to compute the fitness between model and experimental data based on features such as number of spikes, action potential height, action potential width, spike timing, steady state membrane voltage, depolarization and hyperpolarization time constants, and after-hyperpolarization waveform. Channel conductances and membrane properties were varied within a linear range, while kinetic parameters (half activation voltage and time constant) were varied with a multiplicative parameter between 0.5 and 2. Optimizations utilized the Neuroscience Gateway Portal (***Sivagnanam et al., 2013***).

### Calcium dynamics

Intracellular calcium concentration was modeled with diffusion, calcium-binding buffers, and transmembrane calcium pumps. One-dimensional radial diffusion of buffers and calcium was modeled with concentric shells in dendrites, while spine heads and necks implemented one-dimensional axial diffusion in cylindrical slabs connected from the spine neck to the submembrane shell of the dendritic shaft (***Anwar et al., 2014***). Transmembrane calcium extrusion mechanisms—plasma membrane calcium ATPase (PMCA) in every compartment and sodium-calcium exchanger (NCX) limited to spines—were modeled with Michaelis-Menten kinetics. The calcium-binding buffers included calbindin and calmodulin (N and C terminals), which could diffuse between calcium compartments, and an endogenous immobile buffer (representative of several potential biological mechanisms that buffer calcium without diffusing) (***Matthews et al., 2013**; **Matthews and Dietrich, 2015***).

The sources of calcium influx in the model included the voltage gated calcium channels and the NMDAR synaptic channel. Calcium concentration in the submembrane shell was used for calcium-dependent channel activation (BK, SK, CaCC) or inactivation (R-, N-, and L-type calcium channels).

### Synaptic channels

Excitatory NMDAR and AMPAR synaptic channels were included on spine heads, and inhibitory GABA_A_ channels were included on the dendritic shaft. Channels were modeled with dual exponential kinetics, and the NMDA channel included voltage-dependent magnesium blocking.

### Plasticity rule

The calcium based plasticity rule used a dual amplitude and duration threshold for spine calcium concentration to determine LTP and LTD, as we previously described (***Jędrzejewska-Szmek et al., 2017***). For LTD, spine calcium had to exceed the amplitude threshold, ***TA**_D_*, of 0.33 μM for longer than the duration threshold of 28 ms, while for LTP the amplitude threshold, ***TA**_P_*, was 0.53 μM and the duration threshold was 3.3 ms. Once thresholds were exceeded, the synaptic weights were updated at each time step toward their maximum or minimum. For LTP, while thresholds were exceeded, synaptic weight was increased according to:

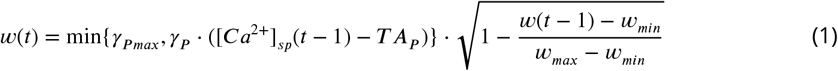

where *γ_P_* is a gain factor for potentiation, *γ_Pmax_* is the maximum allowable gain value, [*Ca^2+^*]_*sp*_ is spine calcium concentration, ***TA**_P_* is the amplitude threshold for potentiation, *w_max_* is maximum allowable synaptic weight (2.0), and *w_min_* is minimum allowable synaptic weight (0). Similarly, when LTD thresholds were exceeded, weight was decreased according to:

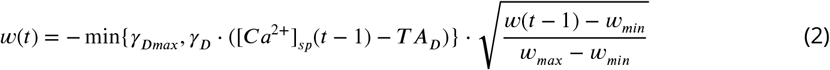

where *γ_D_* and *γ_Dmax_* are the gain and maximum allowable gain for depression, and ***TA**_D_* is the amplitude threshold for depression.

### Synaptic inputs

*In vivo* cortical spike trains were obtained from a CRCNS.org repository, consisting of recordings from 25 simultaneously recorded anterior lateral motor cortex pyramidal neurons with 90 repeated trials (***Li et al., 2015***). A single initial trial of model input consisted of 200 spike trains, which were selected from 22 similar trials (excluding neurons that were inactive within a trial) in order to preserve within-trial correlations between neurons.

To generate controlled trial-to-trial variability of spike timing, the initial trial was repeated 10 times with random jitter of each spike on each repetition. Trial-to-trial variability was limited to standard deviation of spike timing while constraining the same total number of spikes per spike train within each trial. The random jitter was generate from a truncated normal distribution using a standard deviation of 1, 10, 100, or 200 ms (truncated such that no value outside the start time or end time of the trial was selected). Experiments consisted of 10 trials with a single standard deviation. Trials were separated by a 1 second intertrial interval, during which time membrane potential returned to resting potential and spiking activity ceased.

We also implemented a different type of trial-to-trial variability that allows variability of spike rate for individual synapses, but maintains the same spike timing to the neuron as a whole. Variable spike rate was implemented by randomly moving individual spikes from one presynaptic input train to another, with the probability of each spike being moved between 10-100 %. To maintain the distribution of spike counts per trial, the randomly selected target train probability was weighted by the number of spikes of the target trains. This ensured both the overall spike timing pattern to the whole neuron was maintained as well as the distribution of spike counts to synapses, while allowing small variations in spike rate for each synapse from trial to trial.

To demonstrate the robustness of the results to the spatial pattern of synaptic inputs, a single trial was repeated using 1000 different mappings of spike trains to synapses. This preserved the overall instantaneous firing rate to the neuron, and isolated the contribution of spatial patterns.

In addition to excitatory inputs, the neuron also received two types of inhibitory inputs. Inhibitory inputs were constructed from Poisson processes with mean firing rates consistent with both striatal fast spiking interneurons (FSIs) and low threshold spiking interneurons (LTSIs). Inhibitory trains were active for the entire 21 second experiment duration of the 10 repeated trials.

FSI inputs were generated with a mean firing rate of 12 Hz (***Owen et al., 2018***) and targeted densely proximally (within 80 microns of the soma), while LTSI inputs were generated with a mean firing rate of 8 Hz (***Sharott et al., 2012***) and targeted distally (greater than 80 microns from the soma).

### Analysis

Analysis of simulations used Python3 and the following python packages: Numpy, Scipy, Pandas, Scikit-learn, Statsmodels, and Matplotlib. For analysis relating spatiotemporal input patterns to magnitude and direction of synaptic weight change, we introduce a “weight-change triggered average,” which is constructed by grouping synapses per trial into bins based on the value of the weight change following a trial, computing the instantaneous firing rate of the spike train input to each given synapse, and averaging the instantaneous firing rates within each bin of synaptic weight change. This is analogous to a spike-triggered average (***Schwartz et al., 2006***), but using the continuous valued, trial-level weight change instead of the spike.

Random forest regression was applied to the instantaneous firing rate to a synapse, the correlation between synapse firing rate and neighboring synapse firing rate, and the distance from spine to soma. Random forest regression applies a non-linear method for predicting the weight change from a set of features, such as pre-synaptic firing rate. The prediction is determined from a set of hierarchical rules, where each rule partitions the weight change based on a single feature. A non-linear method was required because of the non-linear relationship between weight change and pre-synaptic firing. The instantaneous firing rate of direct input was discretized into 1-5 features (time samples), where 1 time sample calculates the mean firing rate for the entire trial, and 5 time samples calculates mean firing frequency for subsequent 200 ms time intervals. To determine the optimal features for predicting the weight change, a random forest regression was performed for each combination of features. For each regression, the trial-level weight changes were randomly subdivided into a testing set (1/N of the data) and a training set (the remainder of the data); subdividing the data and performing the random forest regression was repeated N times. Then analysis of variance was used to determine which combination of features best predicted the weight change.

