## Supplemental Figures for "Synaptic plasticity is predicted by spatiotemporal firing rate patterns and robust to *in vivo*-like variability"

**Supplementary Figures**

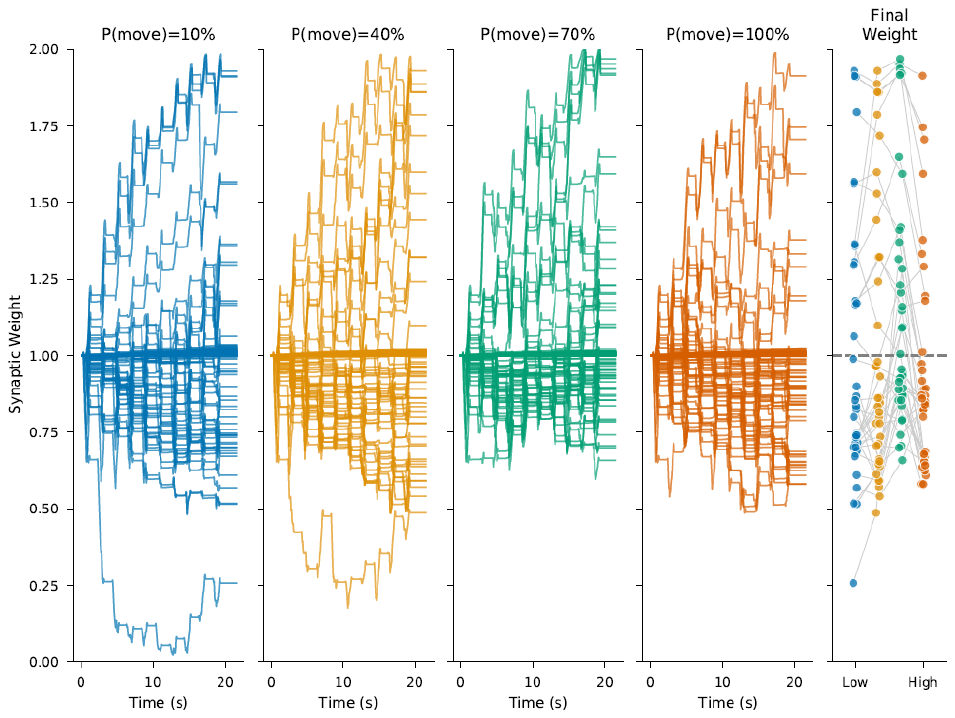
**Figure 4 – supplementary figure 1. Synaptic weight is robust to trial-to-trial variability created by moving spikes between trains.** Individual spikes were moved from one presynaptic input train to another train, with probability between 10 and 100%.

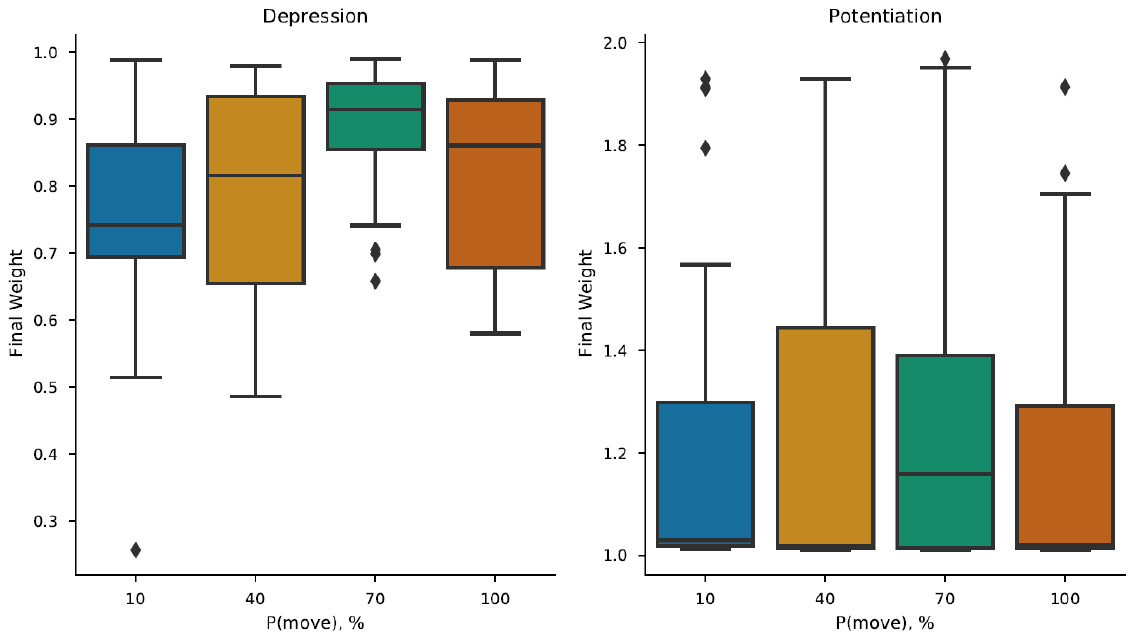
**Figure 4 – supplementary figure 2.** **Distribution of final weights grouped by potentiation and depression for variability introduced by moving spikes between trains.** Correlation of ending synaptic weight versus variability was significant for depressing synapses (R=0.221, p=0.008, N=142 events), but not for potentiating synapses (R=0.014, p=0.873, N=129 events).

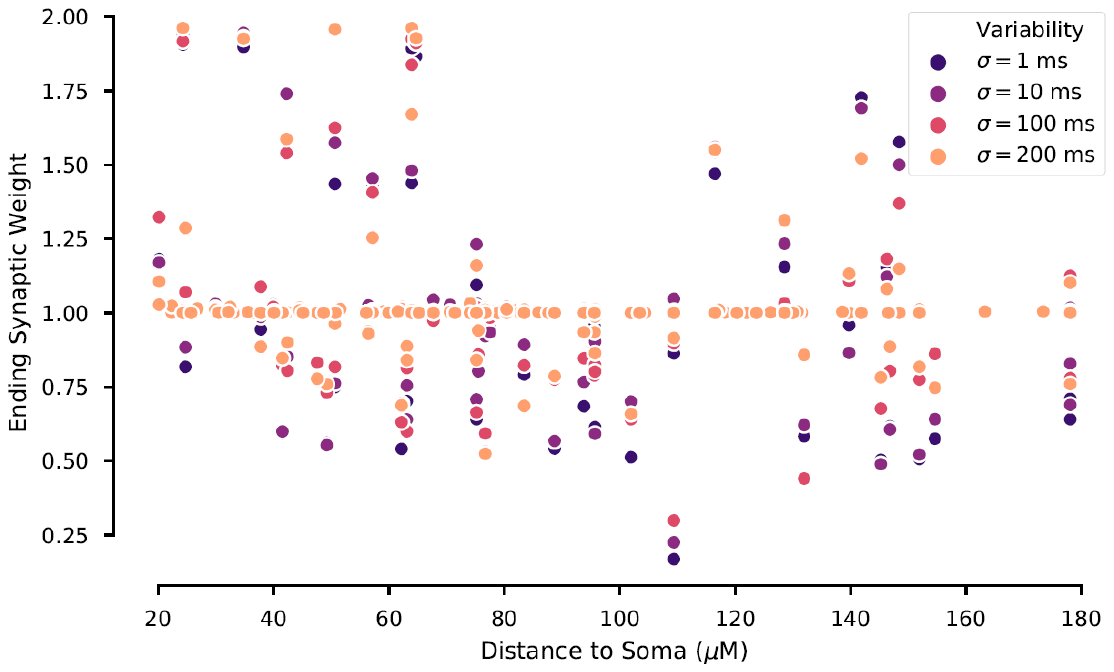
**Figure 5 – supplementary figure 1. Ending synaptic weight versus distance to soma for jittered spike train variability. C**orrelation between ending synaptic weight and distance to soma is not significant for σ= 100 ms (R=-0.23, p=0.07) or 200 ms (R=-0.21, p=0.11), but is significant for σ= 12 ms (R=-0.25, p=0.04) or 10 ms (R=-0.25, p=0.04).

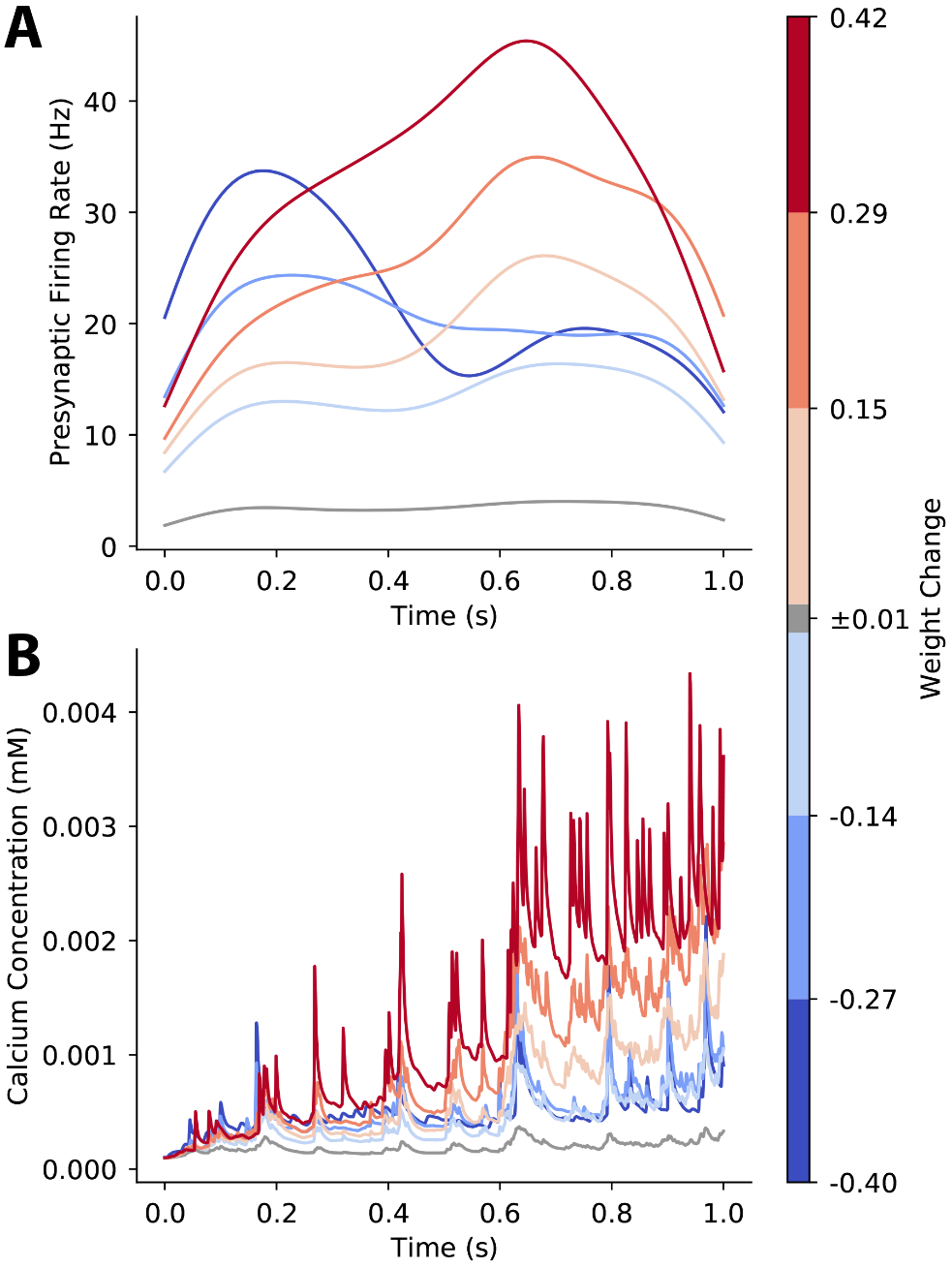
**Figure 6 - supplementary figure 1** – **Weight change triggered average pre-synaptic firing rate and calcium concentration.** Variability is introduced by moving spikes between trains, to maintain the same instantaneous firing rate overall to the neuron. **A.** Synapses that potentiate experience a late peak firing rate. **B.** Size of potentiation correlates with calcium amplitude during the second half of the trial, whereas size of depression correlates with calcium amplitude during the first half of the trial.

**Figure 7 supplementary figure 1. Neighboring synaptic activity when variability is created by moving spikes between trains.** **A.** Weight change triggered synaptic input to neighbors has similar temporal dynamics for synapses that potentiate and synapses that depress, though peak firing is higher for synapses that potentiate. **B.** Correlation between direct and neighboring synapses is high for synapses that potentiate and low or negative for synapses that depress.

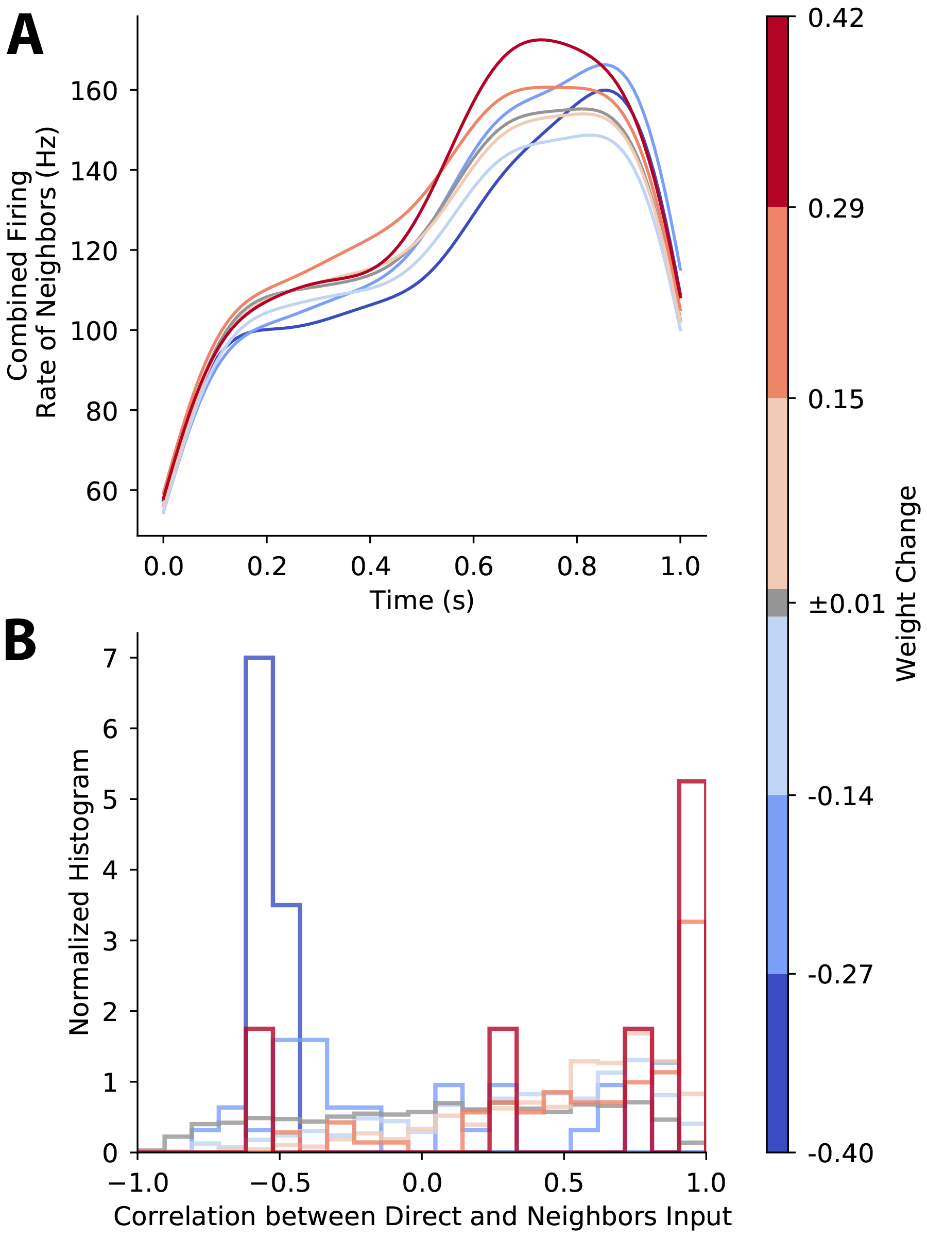

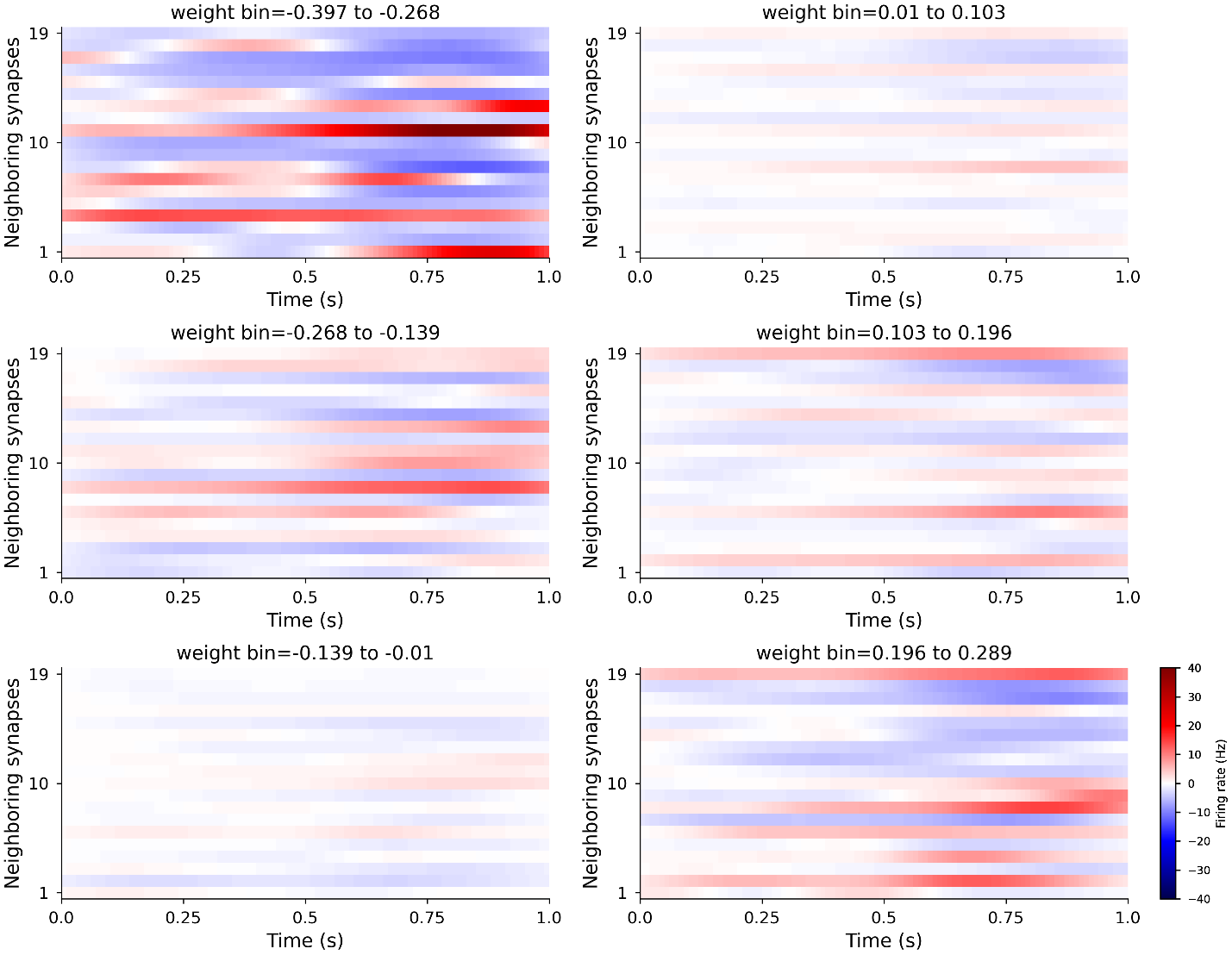
**Figure 7 – supplementary figure 2. Neighboring synaptic activity for 20 nearest neighbors.** Instantaneous firing rate, indicated by color, is relative to mean firing of all synapses. Synapses that strongly potentiate or depress have neighboring synapses with both high and low input firing rates.

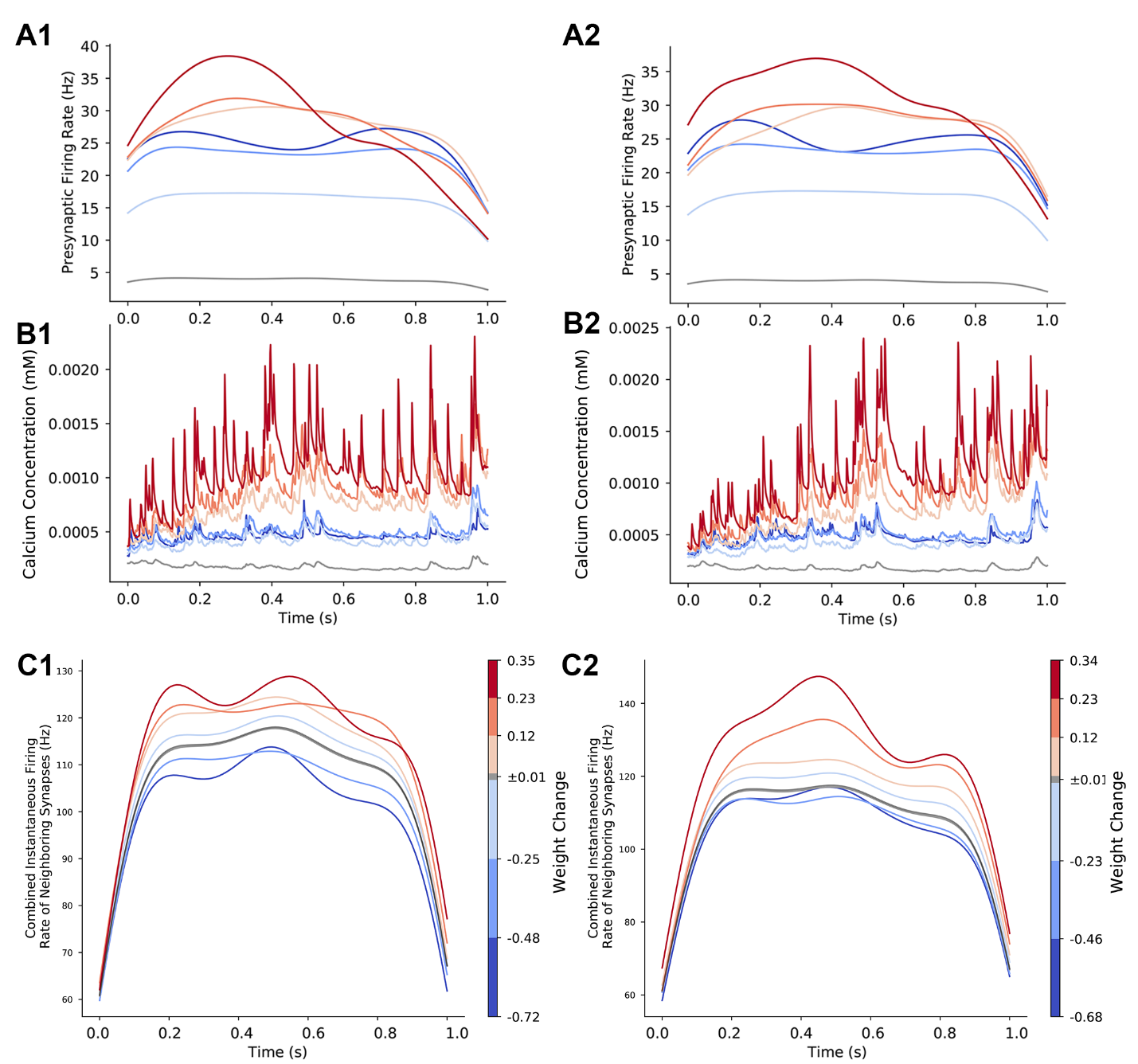
**Figure 7– supplementary figure 3. Temporal pattern of input predicts weight change for different variations of the mapping from spike trains to synapses.** Weight change triggered average pre-synaptic firing rate and calcium concentration for different variations of the mapping from spike trains to synapses. **A.** Weight change triggered pre-synaptic firing rate for two different sets of 200 trials. Synapses that potentiate have a transiently high firing rate, though the time of the firing rate increase varies. **B** Calcium concentration determines direction of plasticity as shown by the weight-change-triggered-average. Calcium concentration for two different sets of 200 trials. Regardless of whether peak synaptic firing rate occurs early or late in the trial, calcium concentration is highest during the second half of the trials. **C** Synapses that potentiate had much higher firing rate than synapses that depressed.

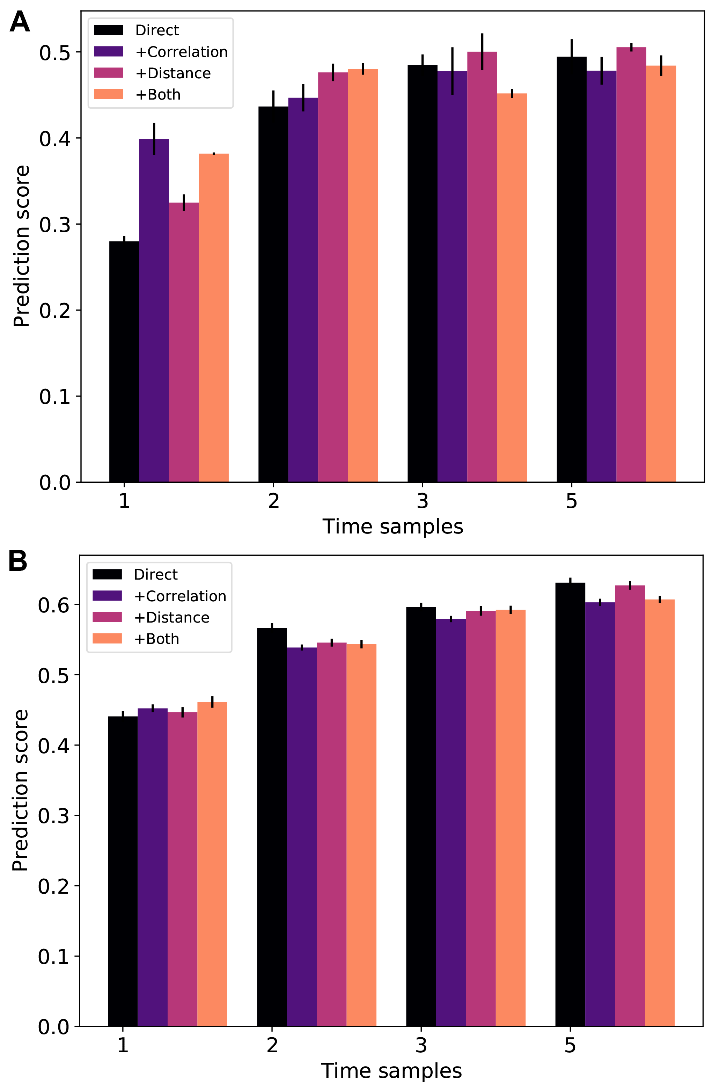
**Figure 8 – supplementary figure 1. Temporal pattern of input predicts weight change better than mean firing rate.** Using distance of synapse to soma (+Distance) or correlation between direct input and input to neighbors (+Correlation) did not improve the weight change prediction. Prediction score is the coefficient of determination, R^2^, of the predicted weight change for the test set. **A.** Variability is created by moving spikes between trains. N=4 regressions for each combination of features. ANOVA shows that increasing number of time samples improves the score (F(3,76)= 64.03, P<0.0001). **B.** Variability is created with different mapping from spike trains to synapses**.** For each set of 200 trials, we performed 4 random forest regressions using 75% of the events in the training set and the remaining 25% of events for testing. N=20 regressions for each combination of features. ANOVA shows that increasing number of time samples improves the score (F(3,396)= 630.9, P<0.0001).
